# Proteomic and Physiological Signatures of Altitude Adaptation in a *Myrsine coriacea* Population under Common Garden Conditions

**DOI:** 10.1101/2023.12.01.569649

**Authors:** Roberta Pena da Paschoa, Vitor Batista Pinto, Jéssica Priscilla Pereira, Paulo Cezar Cavatte, Mário Luís Garbin, Tiago de Oliveira Godinho, Lucas Rodrigues Xavier, Tatiana Tavares Carrijo, Vanildo Silveira

**Affiliations:** Laboratório de Biotecnologia, Centro de Biociências e Biotecnologia (CBB), Universidade Estadual do Norte Fluminense Darcy Ribeiro (UENF), Av. Alberto Lamego, 2000, Campos dos Goytacazes, RJ, 28013-602, Brazil; Unidade de Biologia Integrativa, Setor de Genômica e Proteômica, UENF, Brazil; Universidade Federal do Espírito Santo, Centro de Ciências Exatas, Naturais e da Saúde, Depto. Biologia, Lab. Botânica, Alto Universitário, Guararema, Alegre, ES, Brazil; Reserva Natural Vale, Rodovia BR 101, km 122 s/n Zona Rural, Linhares - ES, 29900-111, Brazil; Laboratório de Biologia Celular e Tecidual, CBB, UENF, Brazil

**Keywords:** Altitude, Altitudinal gradient, Climate change, Comparative proteomics, *Myrsine coriacea*, Natural population, Physiological parameters, Primulaceae

## Abstract

Plants exhibit phenotypic plasticity in response to environmental variations, which can lead to stable genetic and physiological adaptations if exposure to specific conditions is prolonged. *Myrsine coriacea* demonstrates this through its ability to thrive in diverse environments. The objective of the article is to investigate the adaptive responses of *M. coriacea* by cultivating plants from seeds collected at four different altitudes in a common garden experiment. Through integrated physiological and proteomic analyses, we identified 170 differentially accumulated proteins and observed significant physiological differences among the populations. The high-altitude population (POP1) exhibited a unique proteomic profile with significant down-regulation of proteins involved in carbon fixation and energy metabolism, suggesting a potential reduction in photosynthetic efficiency. Physiological analyses showed lower leaf nitrogen content, net CO_2_ assimilation rate, specific leaf area, and relative growth rate in stem height for POP1, alongside higher leaf carbon isotopic composition (δ*13C*) and leaf carbon (*C*) content. These findings provide insight into the complex interplay between proteomic and physiological adaptations in *M. coriace*a and underscore the importance of local adaptations. This study enhances our understanding of how altitude-specific selection pressures can shape plant molecular biology and physiology, offering valuable perspectives for predicting plant responses to global environmental changes.

**Highlight:** This study unveils proteomic and physiological adaptations in a high-altitude *M. coriacea* population with reduced carbon fixation and energy metabolism.

## Introduction

In a rapidly changing climate, plants face distinct environmental challenges compared to those experienced by their ancestors in recent evolutionary history. One way in which plants respond to these changes is through environmentally induced changes in phenotype (Mendonça *et al*., 2019; Pérez-Ramos *et al*., 2019). Plant species can adjust to these new conditions through phenotypic plasticity, adapt through natural selection, or disperse and establish in areas with conditions similar to those where they are adapted, and these options are not mutually exclusive (Crispo, 2008; Malhi *et al*., 2010).

Understanding how plants respond to environmental changes is essential for understanding the processes of plant adaptation and speciation. Altitude gradients are particularly intriguing due to their pronounced changes in various physical environmental characteristics, such as temperature, atmospheric pressure, humidity, sunlight hours, ultraviolet radiation, wind, season duration, and geology (Korner, 2007; Chen *et al*., 2014; Ahmad *et al*., 2016; Rahman *et al*., 2019). Altitude gradients present challenges for successful plant adaptation. As altitude increases, it is observed that certain plant species are replaced by others, giving rise to different vegetation types. Some species are restricted to a specific area, while others grow across a wide range of altitudes, adapting to particular conditions through phenotypic plasticity and genetic modification (Clausen et al., 1940; Hovenden and Vander Schoor, 2004; Byars et al., 2007).

Plant populations occurring along an altitudinal gradient can vary in their morpho-anatomical and physiological characteristics (Ahmad *et al*., 2016; Rahman *et al*., 2020). While physiological variations occurring in plants with increasing altitude are well described (Ahmad *et al*., 2016; Hniličková *et al*., 2016; Ibañez *et al*., 2017; Liu *et al*., 2020; Lim and Burns, 2022), the underlying processes leading to these phenotypic variations are not clearly understood. The continuous challenge lies in identifying the molecular pathways and genes of adaptive significance in this context. Our understanding remains incomplete regarding the specific genes and proteins intricately involved in the local adaptation of diverse populations within tropical species. Consequently, the translation of genetic variations into phenotypic adaptations across varied tropical environments still contains gaps in information. Common garden experiments help in understanding local adaptation by enabling the assessment of the relative importance of plastic and molecular responses within the same species (de Villemereuil *et al*., 2016)

Proteomics can significantly contribute to the elucidation of the molecular mechanisms involved in species adaptation to diverse environments (Diz *et al*., 2012; Li *et al*., 2014; Ma *et al*., 2015; Voelckel *et al*., 2017). A comprehensive understanding of how proteins are regulated in species with a wide distribution range holds the potential to provide valuable insights into the impact of environmental changes on organisms, the evolutionary processes shaping proteins over time, and the strategies employed by organisms to adapt to their specific environmental conditions.

*Myrsine coriacea* is an angiosperm with a broad capacity to inhabit different environments (Sánchez-Tapia *et al*., 2018), exhibiting substantial phenotypic variation (Pipoly, 1991; Ricketson and Pipoly III, 1997; Carrijo *et al*., 2017). In the Atlantic Forest, the species can be found from coastal ecosystems to high-altitude Campos (Carrijo *et al*., 2017). The genetic (Paschoa *et al*., 2018), morphological, and physiological (Pereira *et al*., 2022) diversity in populations distributed across an altitudinal gradient in the Atlantic Forest seems to indicate high phenotypic plasticity, considering the demonstrated capacity of these plants to acclimate to diverse environmental conditions. A better understanding of the influence of the relative roles of genetic and environmental factors on phenotypic variation in *M. coriacea* helps to understand the potentially adaptive phenotypic variation in plants in nature and the adaptive and survival strategies of populations.

In this study, we subjected plants germinated from *M. coriacea* seeds originating from natural populations distributed across an altitudinal gradient to common garden conditions. The objective of the article is to investigate the adaptive responses of *M. coriacea* by cultivating plants from seeds collected at four different altitudes in a common garden experiment. Our findings provide new insights regarding the molecular mechanisms involved in the local adaptation of different populations of *M. coriacea*.

## Materials and Methods

### Seed collection and common garden establishment

The seeds were collected from natural populations located in four municipalities in Espírito Santo State, Brazil, along a 1,517 m altitude gradient (Table 1). Ripe fruits were collected in November 2017 from four branches of ten selected adult trees in each population. After collection, the fruits were subjected to mechanical scarification using a steel sieve and dried in the shade. Then, the seeds were immediately taken to a greenhouse located in Jerônimo Monteiro-ES (20°47’45’’/41°24’18’’). In the nursery, the seeds were planted in polypropylene trays with a capacity of 19.5 cm³ containing washed sand substrate. In March 2018, the seedlings were transplanted into polypropylene tubes with a capacity of 280 cm³ filled with a commercial substrate Tropstrato, HT Hortaliças (Vida Verde^®^, SP, Brazil). The seedlings were kept for 12 months under shade cloth with irradiance levels reduced by 50% and automated micro-sprinkler irrigation activated three times during the daytime period. In March 2019, the seedlings were transferred to three-liter pots containing a substrate composed of a 3:1 ratio of the same commercial substrate and sand, where they remained until the setup of the field experiments.

**Table 1.** Location of the natural populations of *M. coriacea*.

| Population code | Municipality | Locality | Latitude | Longitude | Altitude (M.S.L) |
| --- | --- | --- | --- | --- | --- |
| POP1 | Dores do Rio Preto | Macieira in Caparaó National Park | 20°27'27"S | 41°48'38"W | 2019 |
| POP2 | Domingos Martins | Parque Estadual Pedra Azul | 20°23'38"S | 41°01'36"W | 1229 |
| POP3 | Mimoso do Sul | Pedra dos Pontões | 20°59'35"S | 41°21'08"W | 930 |
| POP4 | Muqui | Private farm property | 20°53'40S | 41°20'12"W | 657 |

A common garden was established in December 2019 in the municipality of Marechal Floriano (Espírito Santo, Brazil) (20°25’38.41“/40°51’29.08”) located at approximately 914 m altitude. The planting was conducted in a randomized block design, with three blocks containing ten plants from each population, totaling 40 plants per block. A total of 87 mixed seedlings were planted in the borders, spaced 2 meters within rows and 3 meters between rows, in a total area of 1.482 m^2^.

Each block consisted of five rows containing eight individuals (two from each population). The seedlings were planted with a spacing of two meters within the rows and three meters between rows. Planting was carried out in pits with dimensions of 30×30×30 cm, where 200 g of superphosphate (19% P) was added 15 days before planting. To ensure the establishment of the seedlings, regular irrigations were performed using manual watering cans. Weed control was carried out through manual weeding and chemical targeted application of the herbicide Roundup Original^®^ (Bayer CropScience, SP, Brazil) Whenever necessary, ant control was carried out using the ant insecticide (Mirex-SD^®^, SP, Brazil)

### Proteomic analysis

#### Protein extraction and digestion

To trace the proteomic profile of the populations, expanded leaves located throughout the canopy were collected from three individuals per population in each block seven months after the common garden was established. The samples collected in the field were stored in sealed paper bags and covered with silica gel. The leaf collection time started at 12:00 PM and finished at 2:00 PM, with an average temperature of 25 °C, relative humidity of 57%, and wind speed of 10 km/h. Subsequently, the samples were wrapped in aluminum foil and stored at −80 °C until proteomic analysis.

For protein extraction, the three individuals collected in each population within each block were pooled and macerated under liquid nitrogen to comprise each biological replicate. protein extraction. Proteins were extracted from 300 mg of powered fresh matter (FM) aliquots using the trichloroacetic acid (TCA)/acetone precipitation method described by Damerval et al., (1986), with modifications as described by Passamani *et al.,*(2017). The protein concentration was determined using the 2-D Quant Kit (Cytiva, Marlborough, MA, USA).

Before the trypsin digestion step, aliquots of 100 µg of proteins were precipitated using the methanol/chloroform methodology to remove any interferents from the samples (Nanjo *et al*., 2012). After precipitation, the samples were resuspended in a 7 M urea/2 M thiourea buffer, and tryptic digestion of proteins (V5111; Promega, Madison, WI, USA; final enzyme-to-protein ratio of 1:100) was performed using the filter-aided sample preparation (FASP) method as described by Wiśniewski et al., (2009) with modifications by Reis *et al*., (2021). The resulting peptides were quantified using the protein and peptide method at 205 nm using a NanoDrop 2000c spectrophotometer (Thermo Fisher Scientific, Waltham, USA).

#### Mass spectrometry analysis

Mass spectrometry was performed using a nanoACQUITY ultra-performance liquid chromatograph (UPLC) coupled to a Q-TOF SYNAPT G2-Si instrument (Waters, Manchester, UK) as described by Reis *et al.,* (2021). The runs consisted of three biological replicates of 2.0 μg of peptides. During separation, the samples were loaded onto a nanoAcquity UPLC M-Class Symmetry C18 5 μm trap column (100 Å, 5 µm, 180 µm × 20 mm, 2D; Waters) at a flow rate of 5 μL/min for 3 min and then onto an analytical reversed-phase nanoAcquity M-Class HSS T3 1.8 μm column (100 Å, 1.8 µm, 75 µm × 150 mm; Waters) at a flow rate of 400 nL/min. The column temperature was set to 45 °C. A binary gradient was used for peptide elution, where mobile phase A consisted of water (Tedia, Fairfield, Ohio, USA) with 0.1% formic acid (Sigma-Aldrich), and mobile phase B consisted of acetonitrile (Sigma-Aldrich) with 0.1% formic acid. The gradient elution was performed as follows: 5% B for 3 min, increasing from 5 to 41% B over 92.00 min, increasing from 41 to 97% B over 96.00 min, holding at 97% B for 100.00 min, and decreasing to 5% B by 102.00 min.

Mass spectrometry was performed in positive mode and resolution mode (mode V), with 35,000 full widths at half maximum (FWHM) and ion mobility, and in data-independent acquisition (DIA) mode. The ion mobility separation (IMS) used an IMS wave velocity of 800 m s^−1^ (HDMS^E^); the transfer collision energy increased from 19 to 55 V in high-energy mode; the cone and capillary voltages were 30 V and 3000 V, respectively; and the source of temperature was 100 °C. For time-of-flight (TOF) parameters, the scan time was set to 0.5 s in continuum mode, and the mass range was 50–2000 Da. Human [Glu1] fibrinopeptide B at 100 fmol µL−1 was used as an external calibrant, and lock mass acquisition was performed every 30 s. Mass spectrum acquisition was performed by MassLynx software (version 4.1, Waters).

### Proteomic data analysis

Spectra processing and database search conditions were performed using ProteinLynx Global SERVER (PLGS) software (version 3.02, Waters). The HDMS^E^ analysis followed the following parameters: Apex3D of 150 counts for low-energy threshold; 50 counts for elevated-energy threshold; 750 counts for intensity threshold; one missed cleavage; minimum fragment ions per peptide equal to three; minimum fragment ions per protein equal to seven; minimum peptides per protein equal to two; fixed modifications of carbamidomethyl (C) and variable modifications of oxidation (M) and phosphoryl (STY); default false discovery rate (FDR) of 1%; automatic peptide and fragment tolerance.

To select the proteome of the species phylogenetically closest to *M. coriacea*, a phylogenetic tree was constructed using the PhyloT website (https://phylot.biobyte.de/) based on the National Center for Biotechnology Information (NCBI) taxonomy. The proteome of the species *Actinidia chinensis* (ID: UP000241394), available on UniProtKB (https://www.uniprot.org/), was used, as it is the species phylogenetically closest to *M. coriacea*.

Label-free quantification analysis was performed using ISOQuant software v.1.8 (Distler *et al*., 2014, 2016). The parameters used were as follows: peptide and protein FDR 1%; sequence length of at least six amino acid residues; and minimum peptide score equal to six. Samples were normalized by a multidimensional normalization process, which corrects peak intensities based on the intensity and retention time domains. The software performed relative protein quantification based on the TOP3 method. Based on the relative abundances of uniquely assigned peptides, the abundances of shared peptides were redistributed to the respective source proteins followed by TOP3-based quantification (Distler *et al*., 2014).

For the comparative analysis, only the proteins present or absent (for unique proteins) in the three biological replicates were accepted for differential abundance analysis. Comparative analyses were performed on POP1 relative to the others. The following comparisons were performed: POP1/POP2, POP1/POP3 and POP1/POP4. Data were analyzed by Student’s t-test (two-tailed). Proteins with a p-value < 0.05 were deemed up-accumulated if the Log_2_ of the fold change (FC) > 0.6 and down-accumulated if the Log_2_ of the FC was less than < −0.6. The mass spectrometry proteomics data have been deposited to the ProteomeXchange Consortium via the PRIDE (Perez-Riverol *et al*., 2022) partner repository with the dataset identifier PXD047424. Functional enrichment analysis, including Gene Ontology (GO) analysis and Kyoto Encyclopedia of Genes and Genomes (KEGG) pathway analysis, was conducted on differentially accumulated proteins (DAPs) using OmicsBox software 3.0.29 (https://www.biobam.com/omicsbox).

The interaction networks of DAPs used the first level of interaction retrieved by STRING version 11.5 (http://string-db.org) search. To generate a protein-protein interaction (PPIs) network, Actinidia chinensis was considered the reference plant species using a minimum required interaction score of 0.7.

To visualize the relationships among the populations, a heatmap plot was generated using the gplots package (Warnes *et al*., 2022) and a principal component analysis (PCA) was performed using the prcomp function in package stats in R (R Core Team, 2023).

### Physiological analysis

Physiological analysis of the populations was performed 13 months after the common garden was established using two plants per population in each block, resulting in 6 biological replicates per population.

For growth analysis, plant height was measured using a graduated ruler (precision of 0.1 cm). Height was determined from ground level to the apical bud of the stem. Based on this measurement, the relative growth rate of stem height (*RGR_H_*, mm.mm^−1^dia^−1^) was calculated with a three-month interval between the two measurements.

To determine the chlorophyll index, four fully expanded leaves located throughout the canopy were selected, and the analysis was performed using ClorofiLOG (CFL 1030, Falker, Rio Grande do Sul, BR). The average value for each plant was used for analysis. Leaf hydraulic conductance (*Kleaf*, mmol m^−2^s^−1^. MPa^−1^) was determined using one leaf per plant following the method described by Martins *et al.,*(2019): *Kleaf* = E* / (Ψw pd - Ψw pm-dark), where E is the transpirational flux and Ψw is the leaf water potential. Leaf vein density (VD; mm mm^_2^) was measured using two leaves per plant, following the protocol outlined by Pereira *et al.,*(2022).

Gas exchange and chlorophyll fluorescence analyses were performed using an infrared gas analyzer (IRGA, Li 6400XT, Li-Cor, Lincoln, USA) equipped with a 2 cm^2^ leaf chamber (Li 6400-40 LCF, Li-Cor, Lincoln, EUA). The measurements of net CO_2_ assimilation (*A,* μmol CO_2_ m^−2^ s^−1^), stomatal conductance (*gs,* mol H_2_O m^−2^ s^−1^) and internal CO_2_ concentration CO_2_ (*Ci*, μmol CO_2_ m^−1^) were conducted between 8:30-10:30 under saturating photosynthetically active radiation of 1.000 µmol m^−2^ s^−1^, a partial pressure of CO_2_ of 400 ppm, and a block temperature of 28 °C.

Ten leaves were collected for specific leaf area (*SLA*; cm^2^ g^−1^) determination and carbon isotopic composition (δ*13C*; ‰) according to Cavatte *et al.,*(2011). Total leaf carbon (*C*) and leaf nitrogen (*N*) concentrations were determined using an elemental analyzer (Carlo Erba, Milan, Italy), while isotopic compositions were measured using a stable isotope ratio mass spectrometer (IRMS Delta Plus, Finnigan Mat, San Jose, USA) at the Center for Nuclear Energy in Agriculture, University of São Paulo. The equations proposed by Farquhar *et al.,*(1989) were used to calculate δ*13C* using the Vienna Pee Dee Belemnite as the standard.

The physiological data were analyzed using a linear model with a source of variation (POP 1 to 4) and one variable at a time (for both physiological variables and isotope measurements). Subsequently, the estimated means of the populations were compared by overlaying confidence intervals (95% confidence intervals), where the lack of overlap indicated significant differences between the means. Finally, the results were presented graphically with bar charts and their respective confidence intervals. All analyses were performed using R (R core Team, 2023), using the function emmeans from the package emmeans (Russell V. Lenth, 2023).

## Results

### Proteomic analysis

The comparative proteomic analysis allowed the identification of 590 proteins in the *M. coriacea* samples from the four populations included in the experimental planting. Comparative analysis between POP1 and the other populations revealed that 170 DAPs (Supplementary Table S1). Heatmap analysis, using the unweighted pair group method using arithmetic averages (UPGMA) algorithm based on the total ion count of proteins in the biological replicates of POP1, POP2, POP3, and POP4, allowed visualization of the separation into two major groups, one comprising POP1 and the other consisting of the remaining populations. This analysis revealed that POP1 exhibited a distinct protein profile, with most proteins displaying lower total ion count values (Fig. 1A).

**Fig. 1.**
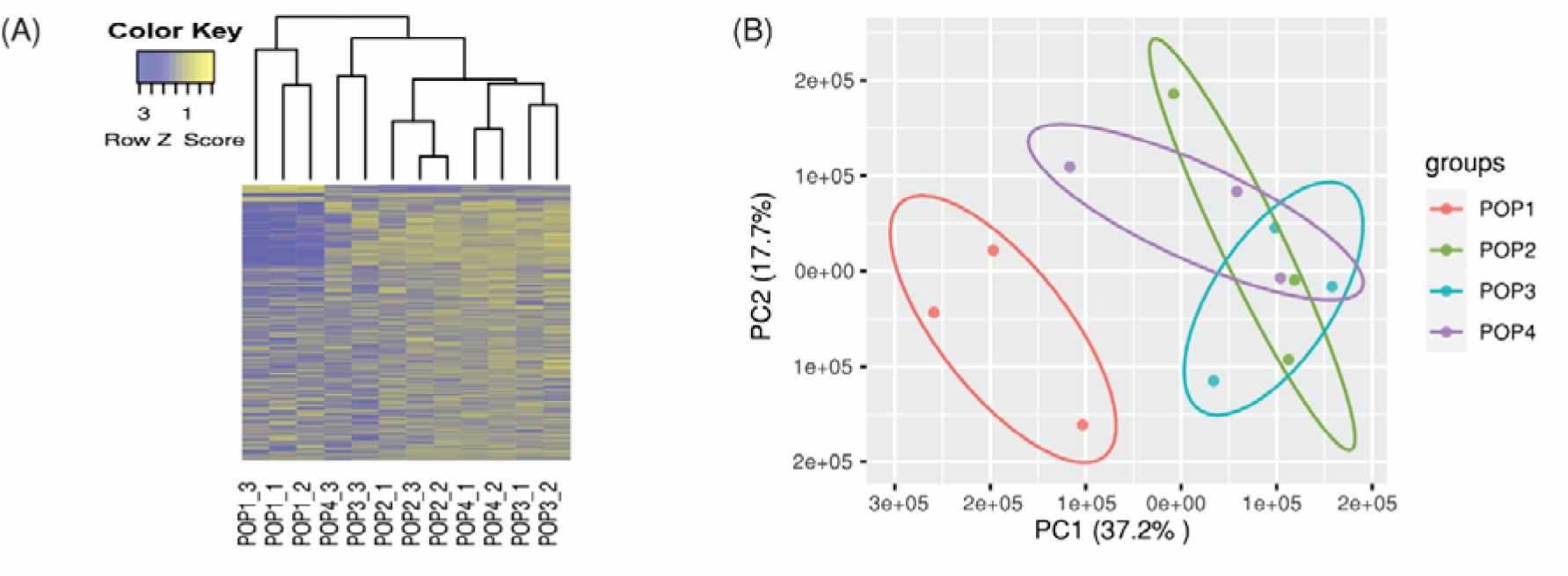
Proteomic analysis relationships between natural populations of *M. coriacea* based on the protein identification and quantitation results. A) Heatmap using the Unweighted Pair Group Method using Arithmetic averages (UPGMA); B) PCA.

Additionally, POP1 distinguished itself from the other studied populations in the PCA analysis (Fig. 1B). Each biological replicate for a specific population was evaluated, and their relationship is indicated by an enclosed circle. The first two eigenvalues of PCA explained 54.9% of the total variation, with 37.2% explained by component 1 and 17.7% by component 2.

In Gene Ontology (GO) analysis, the most representative biological processes in all populations were the “cellular process,” “biosynthetic process,” “response to cadmium ion,” and “generation of precursor metabolites and energy.” Many proteins were found to be down-accumulated in POP1 (Supplementary Fig. S1).

The KEGG pathway analysis revealed that the most representative pathway in all three comparisons was “Carbon fixation in photosynthetic organisms,” followed by “glycolysis/gluconeogenesis” for the POP1/POP2 and POP1/POP3 comparisons and “cysteine and methionine metabolism” for the POP1/POP4 comparison (Supplementary Fig. S2).

The predicted analysis of protein-protein interactions (PPIs) resulted in interactions among potential key proteins involved in the differential regulation of POP1/POP2 (Fig. 2), POP1/POP3 (Fig. 3), and POP1/POP4 (Fig. 4) comparisons.

**Fig. 2.**
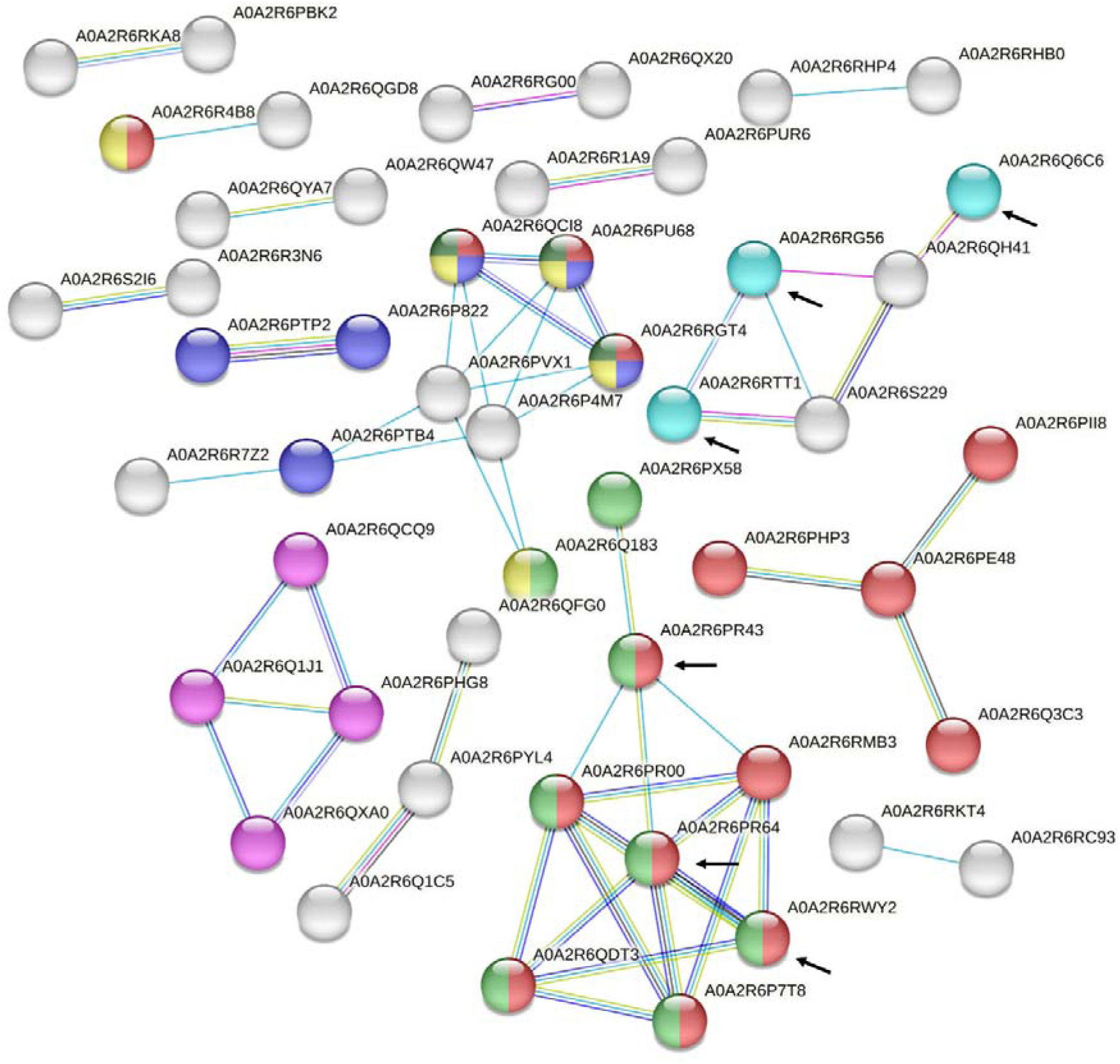
Protein-protein interaction (PPIs) network among the DAPs in the comparative proteomic analysis of POP1/POP2. The red nodes represent proteins involved in “carbon fixation in photosynthetic organisms,” the blue nodes are related to “glyoxylate and dicarboxylate metabolism,” the light green nodes are involved in “glycolysis/gluconeogenesis,” the maroon nodes are related to “pyruvate metabolism,” the dark green nodes represent proteins involved in “cysteine and methionine metabolism,” the pink nodes are involved in “amino sugar and nucleotide sugar metabolism,” and the light blue nodes are involved in “photosynthesis.” The colored lines indicate various types of interaction evidence (dark blue: co-occurrence; black: co-expression; light blue: database evidence; green: neighboring genes; pink: experimental evidence; red: fusion evidence; yellow: text mining evidence; purple: protein homology). Arrows indicate the proteins highlighted for discussion.

**Fig. 3.**
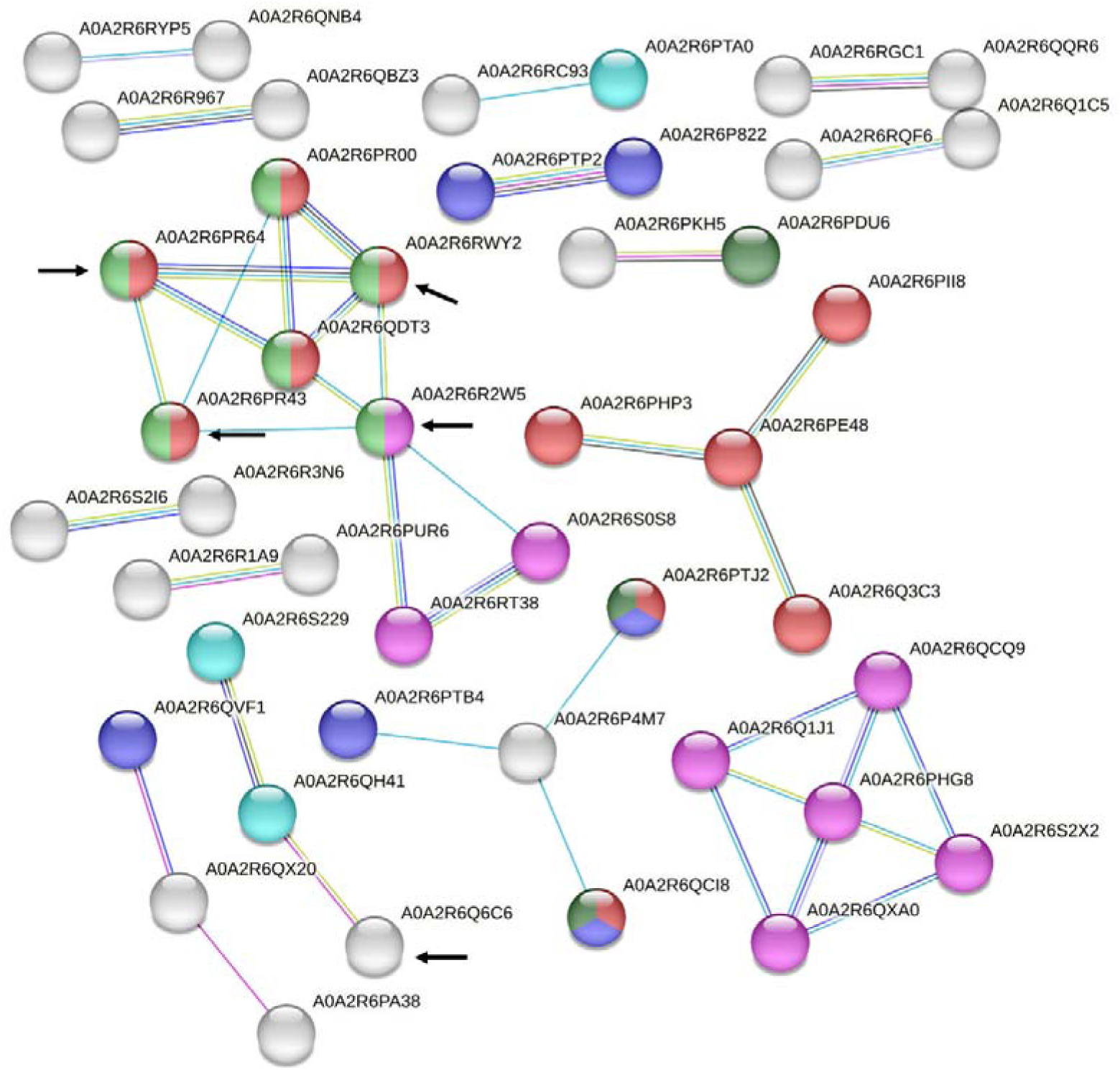
Protein-protein interaction (PPIs) network among the DAPs in the comparative proteomic analysis of POP1/POP3. The red nodes represent proteins involved in “carbon fixation in photosynthetic organisms,” the blue nodes are related to “glyoxylate and dicarboxylate metabolism,” the light green nodes are involved in “glycolysis/gluconeogenesis,” the maroon nodes are related to “pyruvate metabolism,” the dark green nodes represent proteins involved in “cysteine and methionine metabolism,” the pink nodes are involved in “amino sugar and nucleotide sugar metabolism,” and the light blue nodes are involved in “photosynthesis.” The colored lines indicate various types of interaction evidence (dark blue: co-occurrence; black: co-expression; light blue: database evidence; green: neighboring genes; pink: experimental evidence; red: fusion evidence; yellow: text mining evidence; purple: protein homology). Arrows indicate the proteins highlighted for discussion.

**Fig. 4.**
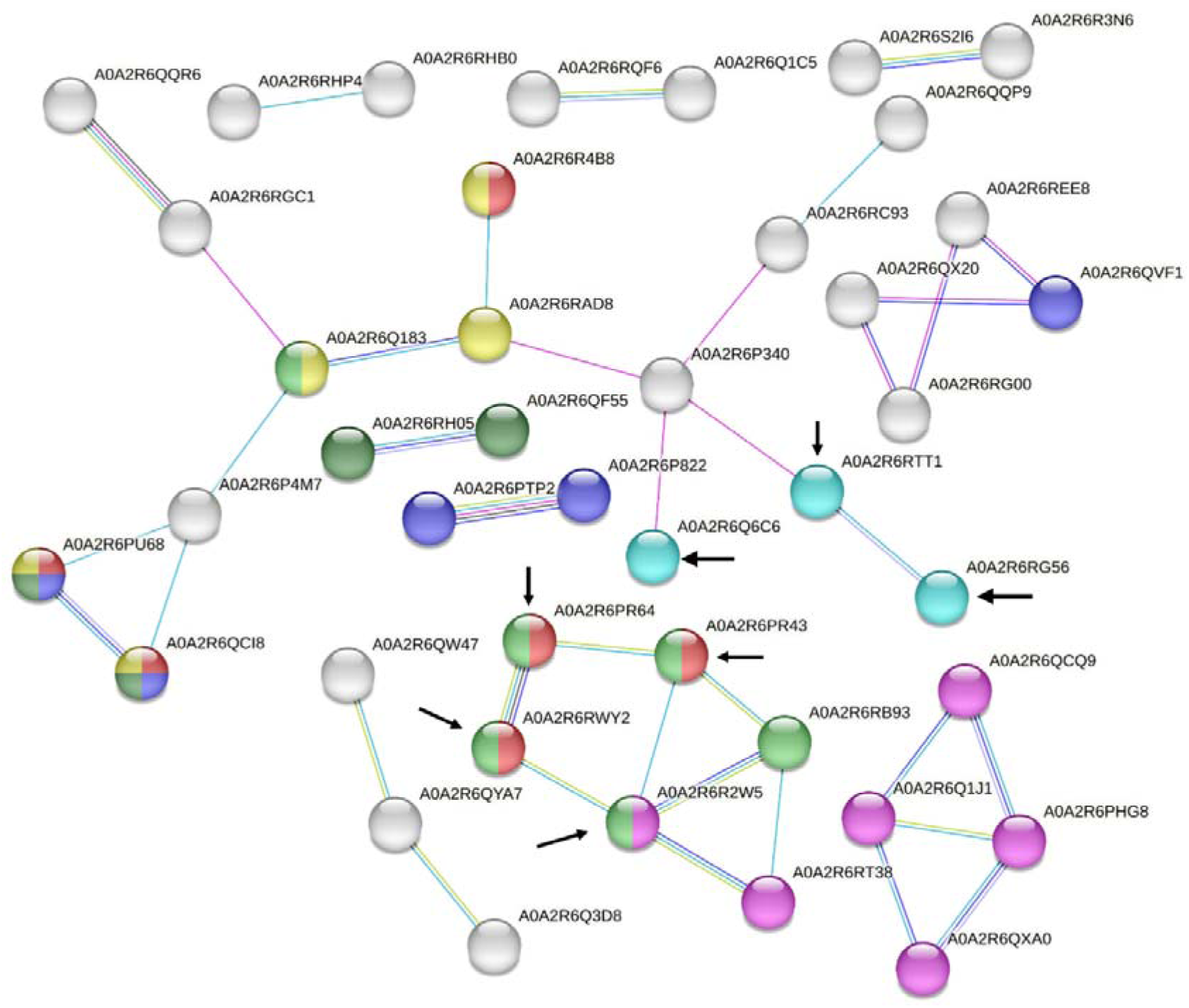
Protein-protein interaction (PPIs) network among the DAPs in the comparative proteomic analysis of POP1/POP4. The red nodes represent proteins involved in “carbon fixation in photosynthetic organisms,” the blue nodes are related to “glyoxylate and dicarboxylate metabolism,” the light green nodes are involved in “glycolysis/gluconeogenesis,” the dark green nodes represent proteins involved in “cysteine and methionine metabolism,” the pink nodes are involved in “amino sugar and nucleotide sugar metabolism,” and the light blue nodes are involved in “photosynthesis.” The colored lines indicate various types of interaction evidence (dark blue: co-occurrence; black: co-expression; light blue: database evidence; green: neighboring genes; pink: experimental evidence; red: fusion evidence; yellow: text mining evidence; purple: protein homology). Arrows indicate the proteins highlighted for discussion.

PPIs among Triosephosphate isomerase (A0A2R6RWY2), Glucose-6-phosphate isomerase (A0A2R6R2W5), Fructose-bisphosphate aldolase (A0A2R6PR43), and Phosphoglycerate kinase (A0A2R6PR64) were visualized in the comparisons between populations (Fig. 2, 3, and 4; highlighted arrows). These proteins are involved in the pathways of “Carbon fixation in photosynthetic organisms” and “Glycolysis/gluconeogenesis” and are down-accumulated in POP1 compared to the other populations. There was the exception of the POP1/POP2 comparison, which was unchanged, but also showed a lower accumulation in POP1 in this comparison. Chlorophyll a-b binding proteins (A0A2R6Q6C6, A0A2R6RTT1, A0A2R6RG56) that were also highlighted in the interaction network, participate in the “photosynthesis-antenna proteins” pathway. These proteins are up-accumulated in POP1 compared to the other populations (Fig. 2, 3, and 4; highlighted arrows).

### Physiological parameters

The analysis of physiological parameters revealed significant differences among the populations (Fig. 5, 6 and Supplementary Fig. S3). δ*13C* (Fig. 5A) and *C* (Fig. 5B) were significantly different in POP1 compared to the other populations, with POP1 showing higher values. POP1 exhibited the lowest values of *N* (Fig. 5C) and *RGR_H_*(Fig. 5D) compared to POP2 and POP4. In *Kleaf* (Fig. 6A), *A* (Fig. 6B), and *SLA* (Fig. 6C), there was also a significant difference between POP1 and POP2, with POP1 showing lower values. There were no significant differences among all four populations for *gs*, *CIT*, *VD,* and *Ci* (Supplementary Fig. S3). POP2, POP3, and POP4 were statistically similar in all physiological analyses.

**Fig. 5.**
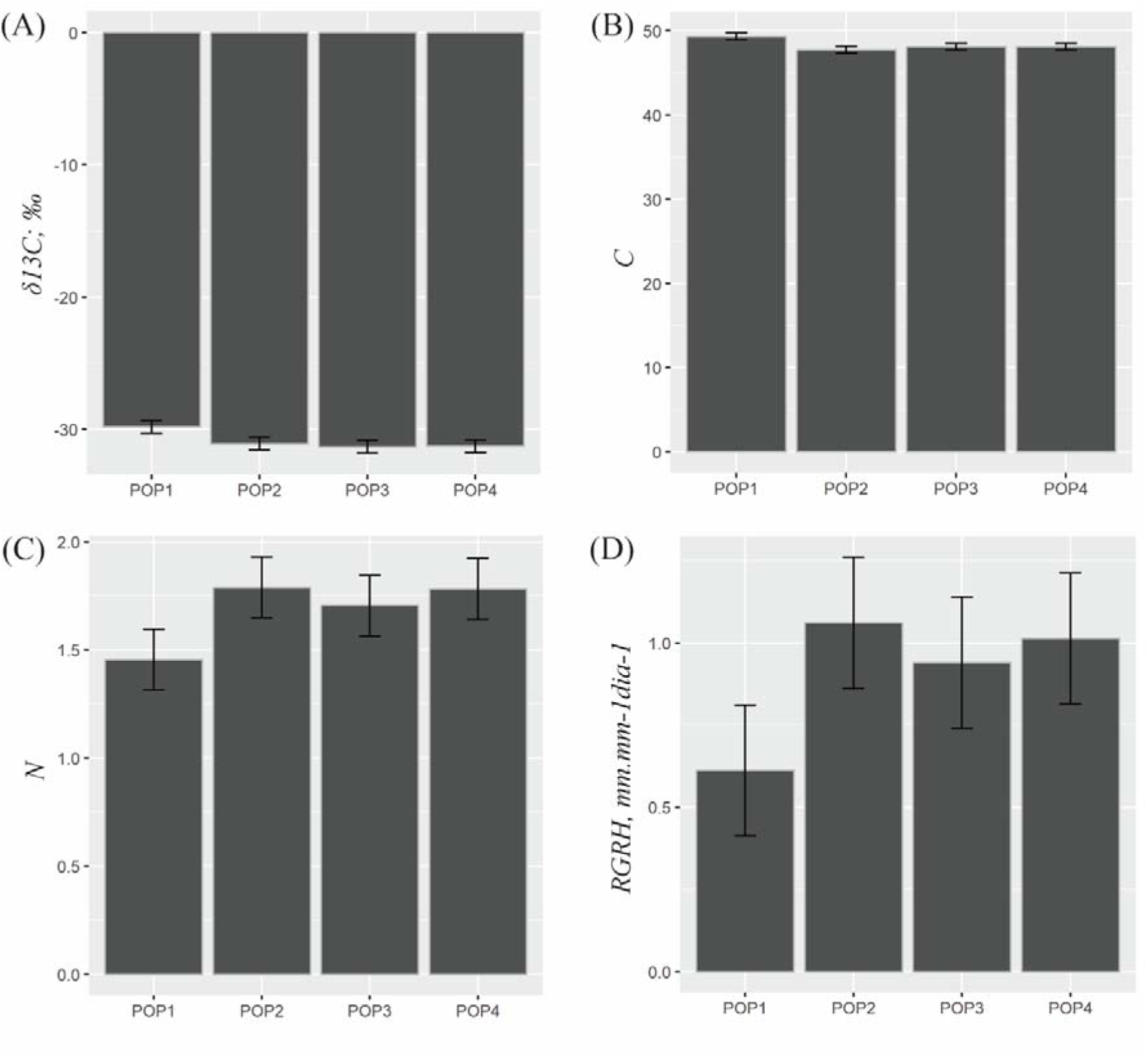
Physiological parameters among the natural populations of *M. Coriacea*. A) Leaf carbon isotopic composition (δ*13C*); B) leaf carbon (*C*); C) leaf nitrogen (*N*); and D) relative growth rate in height (*RGR_H_*). Data are presented as the mean ± CI (95% confidence interval) of six independent replicates The error bars represent confidence intervals.

**Fig. 6.**
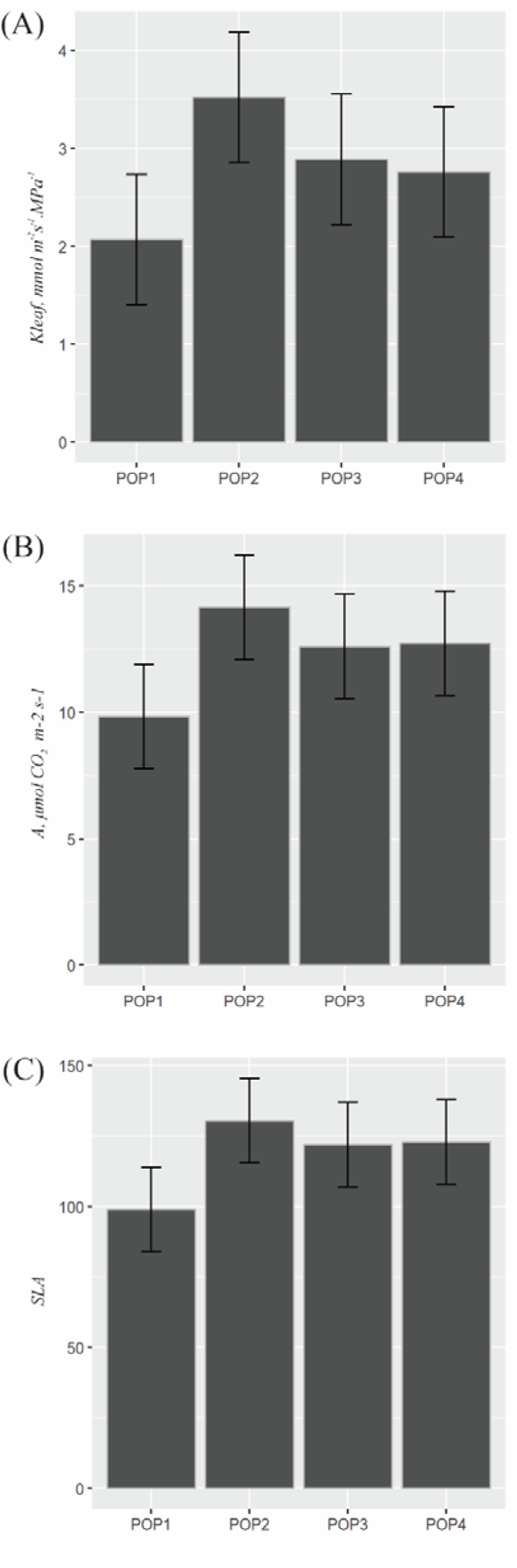
Physiological parameters among the natural populations of *M. Coriacea*. A) Leaf hydraulic conductivity (*Kleaf*); B) net assimilation rate of CO_2_ (*A*); and C) specific leaf area (*SLA*). Data are presented as the mean ± CI (95% confidence interval) of six independent replicates. The error bars represent confidence intervals.

## Discussion

Proteomic analysis revealed differential regulation of proteins among the four populations originating from the altitudinal gradient and subjected to a common garden condition. Based on the cluster analysis with a heatmap and PCA (Fig. 1) of the identified proteins, it was possible to separate the natural populations into two main groups, one containing the highest altitude population (POP1) and the other comprising the remaining populations at lower altitudes. Our results also revealed a differential pattern for POP1 based on physiological analyses (Fig. 5 and 6). These results indicate that the high-altitude population (POP1) (Table 1), exhibits distinct molecular and physiological adaptations when compared to the other populations. This distinction suggests a specific local adaptation of POP1 to the environmental conditions unique to its original high-altitude habitat, which becomes particularly evident when these populations are subjected to common garden conditions at 914 m altitude.

POP1 was separated from the other populations, likely due to the edaphoclimatic differences to which the population is exposed in its original environment. According to the latest update of the Köppen and Geiger climate classification by Alvares *et al.,*(2013), the region of POP1 is characterized as a tropical highland climate, situated at approximately 2019 m altitude, where temperatures can drop to −4 °C in winter. In contrast, the regions of origin of the other populations, POP2 and POP3, are classified as having a warm temperate climate, while POP4 originates from a rainy tropical climate. These latter regions, in comparison to POP1, are marked by lower altitudes and higher temperatures year-round.

Our comparative proteomic analysis revealed that ‘carbon fixation in photosynthetic organisms’ and ‘glycolysis/gluconeogenesis’ were the most representative pathways in POP1, in comparison to the other populations. This was evidenced by a notable down-accumulation of many related proteins (Supplementary Fig. S2; Supplementary Table S1). These results suggest that POP1 exhibits differential protein regulation, particularly in photosynthesis pathways, when subjected to common garden conditions. Natural populations of *M. coriacea* in their native environment showed differences in morphological and stomatal characteristics due to altitude, affecting the photosynthetic capacity of the species at higher altitudes (Pereira *et al*., 2022). Individuals from the high-altitude population (POP1; 2019 m) had leaves with higher vein density (*VD*), trichome density, and leaf carbon isotopic composition (δ*13C*) but lower guard cell length, maximum stomatal conductance, and specific leaf area (*SLA*) compared to the other populations. In our study, δ*13C* also exhibited higher values, and *SLA* showed lower values in POP1 than in the other populations under common garden conditions. However, *VD* did not differ among populations. These results suggest that the species *M. coriacea* is capable of local adaptation in response to specific environmental conditions, such as altitude, through changes in morphological and stomatal characteristics. The high-altitude population (POP1) may have developed specific adaptations, such as increased trichome density, higher δ*13C* levels, and lower *SLA*, to cope with the conditions of high altitude. These adaptations could affect the species’ photosynthetic capacity of the at higher altitudes.

Additionally, alongside the higher values of δ*13C* and *C* content, POP1 exhibited lower *Kleaf* and *A*), suggesting that plants from this population may have reduced their stomatal opening to minimize water loss. δ*13C* is an important indicator of the environmental conditions in which the plant has grown. Soil water availability and climatic conditions are factors that affect this composition, with drought leading to a higher proportion of heavy carbon isotopes in the leaf (Vantyghem *et al*., 2022). This is because the plant tends to reduce stomatal opening to avoid excessive water loss, which can result in lower leaf hydraulic conductance (*Kleaf*) (Jákli *et al*., 2016). Additionally, drought can lead to a reduction in the net assimilation rate of CO_2_ (*A*) (Fresneau *et al*., 2007) and an increased internal concentration of CO_2_ (*Ci*) in the leaf (Nunes *et al*., 1968), indicating lower photosynthetic efficiency in terms of carbon fixation.

The down-accumulation of proteins related to energy metabolism might negatively impact the photosynthetic process in the high-altitude population POP1, when compared to the other populations. In a controlled greenhouse experiment with *Quercus ilex* from different altitude ranges, a differential proteomic profile among populations was revealed, with the high-altitude population exhibiting a down-regulation of proteins related to energy metabolism (San-Eufrasio *et al*., 2021). In our study, we identified down-accumulated proteins in POP1, when compared to the other populations, that were involved in energy metabolism, such as Triosephosphate isomerase (A0A2R6RWY2), Glucose-6-phosphate isomerase (A0A2R6R2W5), Fructose-bisphosphate aldolase (A0A2R6PR43), and Phosphoglycerate kinase (A0A2R6PR64). These proteins were found to participate in predicted protein-protein interactions in the performed comparisons (Fig. 2, 3, and 4).

The decrease in proteins involved in energy metabolism suggests that POP1 has a different energy yield compared to the other populations, consequently reducing its photosynthetic capacity. This hypothesis is consistent with our physiological analysis results in the studied populations. Physiological analyses revealed that POP1 exhibited higher values of *C* and δ*13CI*, and lower values of *A* and relative stem height growth rate (*RGR_H_*). Increasing temperature leads to an increase in the carboxylation rate and a decrease in the specificity of RUBISCO (ribulose-1,5-bisphosphate carboxylase/oxygenase) for CO_2_ (Brooks and Farquhar, 1985). It has been demonstrated that leaves with higher δ*13C* values exhibit lower rates of photosynthesis and stomatal conductance than those with lower values (Vitoria *et al*., 2016). Thus, photosynthesis and growth are affected by temperature, which limits biochemical processes (Cooper and Taiton, 1968).

Nitrogen assimilation is a highly energy-demanding process (Bloom *et al*., 1992). Once absorbed, nitrogen can be assimilated in the root itself or transported to the leaves, where assimilation takes place (Lee *et al*., 1992). Since POP1 exhibited down-regulation of proteins involved in energy metabolism, indicative of different energy yields compared to the other populations and a reduced photosynthetic capacity, it may also interfere with nitrogen assimilation, as evidenced by a lower leaf nitrogen concentration (*N*) in POP1 (Fig. 5C).

Energy deficit is a primary symptom of plants under stress (Baena-González *et al*., 2007). In this study, an up-regulation of Chlorophyll a-b binding proteins (A0A2R6RG56, A0A2R6RTT1, A0A2R6Q6C6) was identified in POP1, when compared to the other populations. The expression of Chlorophyll a-b binding protein encoding genes is affected by light intensity, low temperature, high salinity, drought, and disease (Humbeck and Krupinska, 2003; Caffarri *et al*., 2005; Umate, 2010; Xia *et al*., 2012; Wang *et al*., 2017). Regulating the levels of Chlorophyll a-b binding proteins allows chloroplasts to respond flexibly and rapidly to environmental stresses (Li *et al*., 2020). The composition of pigments is altered to adapt to specific environmental conditions and when a plant is exposed to lower irradiation, it produces more chlorophyll to absorb light and capture more energy (Arena *et al*., 2020). The up-regulation of these Chlorophyll a-b binding proteins in POP1, may be related to the plants’ need to respond to environmental stress, such as lower irradiation and higher temperature compared to their original location.

## Conclusion

The populations studied displayed distinct proteomic and physiological profiles, and the high-altitude population (POP1) demonstrating particularly unique characteristics. In POP1, a notable decrease in proteins involved in energy metabolism was observed, including Triosephosphate isomerase (A0A2R6RWY2), Glucose-6-phosphate isomerase (A0A2R6R2W5), Fructose-bisphosphate aldolase (A0A2R6PR43), and Phosphoglycerate kinase (A0A2R6PR64). This population also showed lower concentrations of *N*, *A*, *SLA* and *RGR_H_*, coupled with higher concentrations of δ*13C* and *C*, indicating a reduced photosynthetic capacity in POP1 under common garden conditions.

Furthermore, the up-regulation of Chlorophyll a-b binding proteins (A0A2R6RG56, A0A2R6RTT1, A0A2R6Q6C6) in POP1 suggests an adaptation to environmental stress, which is significant in the acclimation process of high-altitude populations of *M. coriacea*. The distinct local adaptations in traits such as *N*, *A*, *SLA*, *RGR_H_*, δ*13C*, and *C*, and the observed differentially accumulated proteins related to energy metabolism underscore the role of abiotic selection pressures in the high-altitude environment in driving the proteomic and physiological changes observed in POP1, even under common garden conditions.

These findings highlight the importance of understanding local adaptations in response to abiotic stresses to better predict how *M. coriacea* populations might respond to climate change. Such knowledge is crucial for predicting their survival and genetic diversity in changing environmental conditions.

## Abbreviation list

*A*: Net assimilation rate of CO_2_
*C*: Leaf carbon
*Ci*: Internal CO_2_ concentration
*CIT*: Total chlorophyll
DAPs: differentially accumulated proteins
FASP: filter-aided sample preparation
GO: Gene ontology
*gs*: Stomatal conductance
KEGG: Kyoto Encyclopedia of Genes and Genomes
*Kleaf*: Leaf hydraulic conductivity
*N*: Leaf nitrogen
PCA: principal component analysis
POP1: Population originating from Dores do Rio Preto, at an altitude of 2019 meters
POP2: Population originating from Domingos Martins at an altitude of 1229 meters
POP3: Population originating from Mimoso do Sul at an altitude of 930 meters
POP4: Population originating from Muqui at an altitude of 657 meters
PPIs: protein-protein interaction
*RGR_H_*: Relative growth rate in height
*SLA*: Specific leaf area
TCA: trichloroacetic acid
UPGMA: Unweighted Pair Group Method using Arithmetic averages
*VD*: Vein density
δ*13C*: Leaf carbon isotopic composition

## Acknowledgements

RPP is grateful for the scholarship funding provided by the Coordenação de Aperfeiçoamento de Pessoal de Nível Superior (CAPES), and RLX acknowledges the scholarship funding from the Fundação de Amparo à Pesquisa do Estado do Rio de Janeiro (FAPERJ). We extend our sincere thanks to the Laboratório de Botânica at the Universidade Federal do Espírito Santo for their invaluable assistance with seed collection, physiological analysis support, and the establishment of the common garden.

## Author contributions

RPP, TCC, PCC, and VS conceived and coordinated the research. RPP performed data collection, analysis, and interpretation and wrote the draft of the manuscript. VBP performed data collection and analysis. JPP performed data collection and analysis. PCC, MLG, TOG, and TTC set up the field experimental design. LRX assisted in data analysis and interpretation. All authors revised and edited the final version of the manuscript.

## Data availability

The mass spectrometry proteomics data have been deposited to the ProteomeXchange Consortium via the PRIDE partner repository with the dataset identifier PXD047424. The list of all identified proteins is available in the supplementary material.

