## Supplementary material for "Proteomic and Physiological Signatures of Altitude Adaptation in a *Myrsine coriacea* Population under Common Garden Conditions": Figures S1-S3: Supplementary_Fig_S1-S3.pdf

**Supplementary figure**

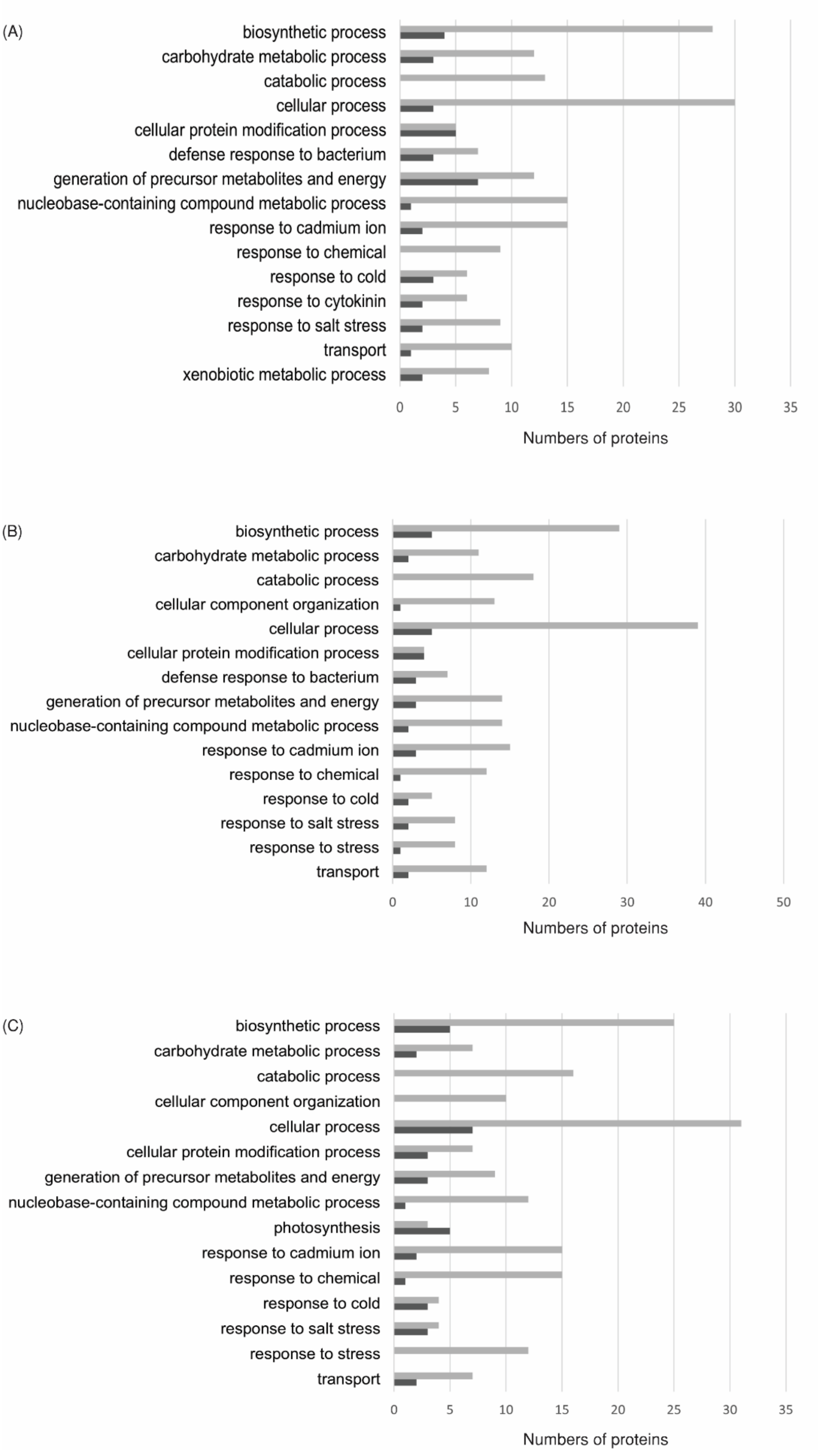


**Supplementary Fig. S1.** Functional classification based on Gene ontology (GO) of DAPs during comparative proteomic analysis between natural populations of *Myrsine coriacea*. A) POP1/POP2; B) POP1/POP3; and POP1/POP4 comparisons. The dark gray bars correspond to up-accumulated proteins, while the light gray bars represent down-accumulated proteins.

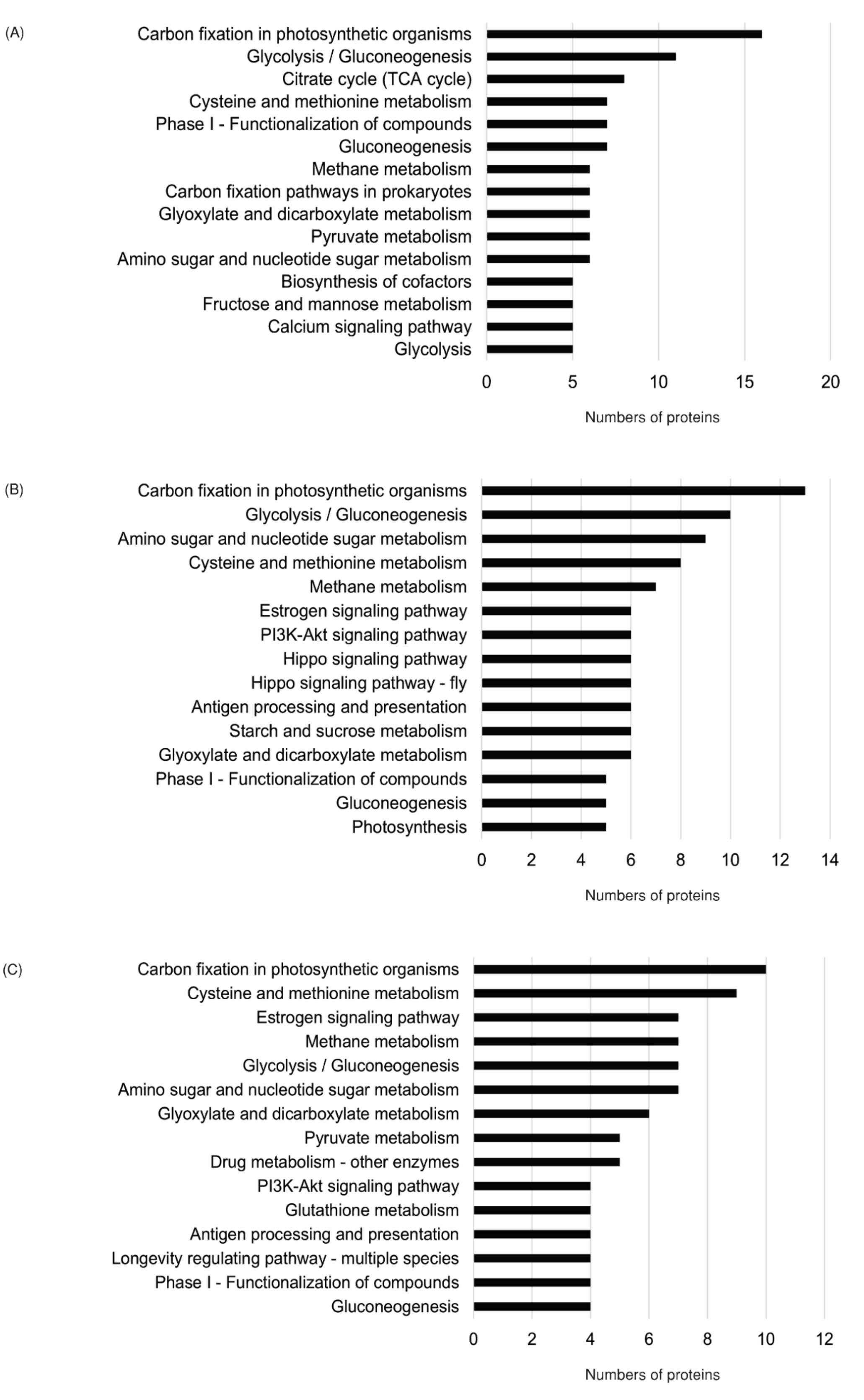


**Supplementary Fig. S2.** KEGG pathways analysis of DAPs during comparative proteomic analysis between natural populations of *Myrsine coriacea*. A) POP1/POP2; B) POP1/POP3; and POP1/POP4 comparisons. The dark gray bars correspond to up-accumulated proteins, while the light gray bars represent down-accumulated proteins.

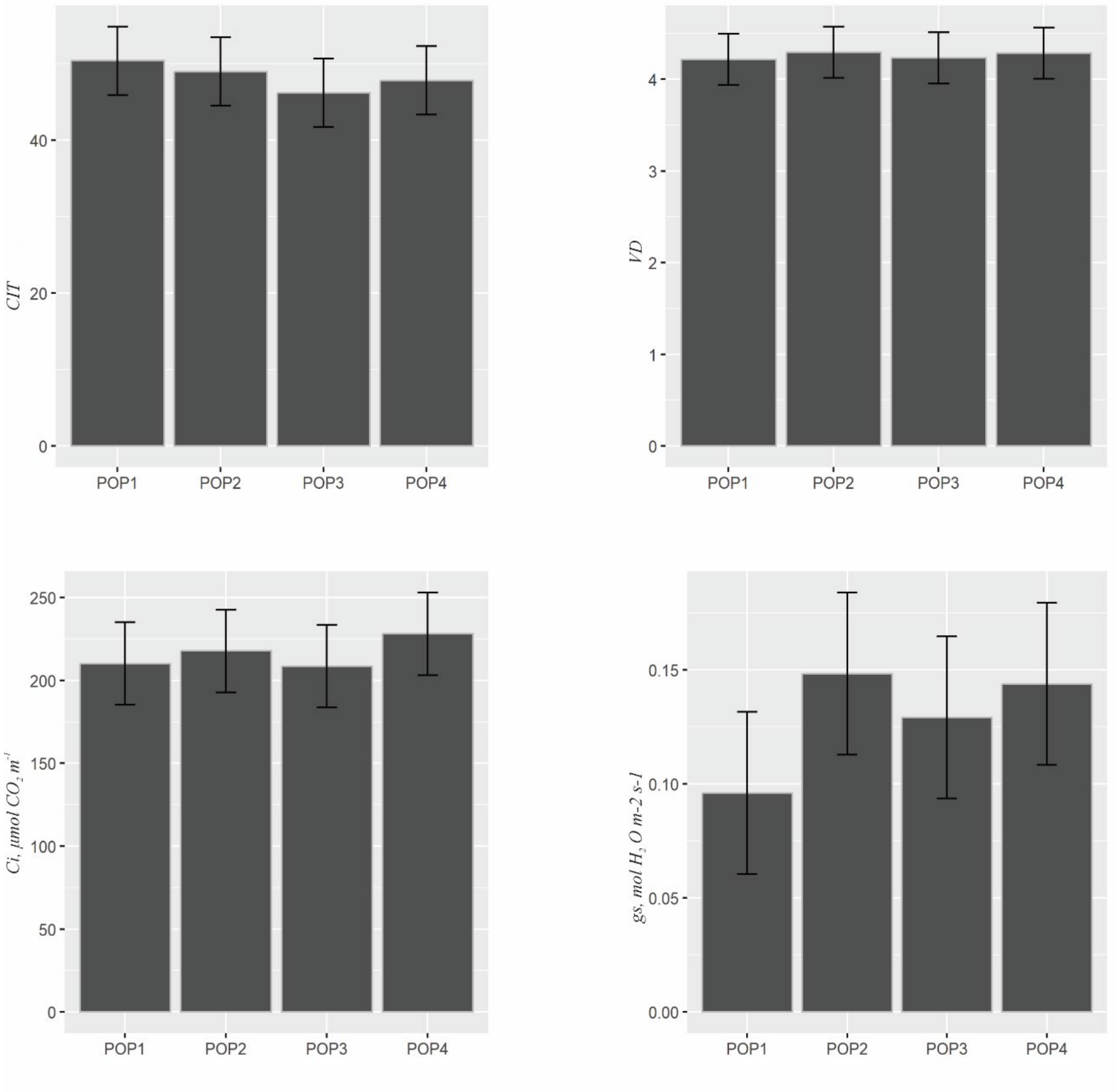


**Supplementary Fig. S3.** Physiological parameters among the natural populations of *Myrsine Coriacea*. A) total chlorophyll (*CIT*); B) vein density (*VD*); C) internal $CO_2$ concentration (*Ci*), and D) stomatal conductance (*gs*). Data are presented as the mean ± CI (95% confidence interval) of six independent replicates. The error bars represent confidence intervals.
